# FACS-based isolation and RNA extraction of Secondary Cells from the Drosophila male Accessory Gland

**DOI:** 10.1101/630335

**Authors:** Clément Immarigeon, François Karch, Robert K. Maeda

**Author notes:** Email addresses of co-authors: François KARCH, Robert K. MAEDA. Corresponding author: Clément IMMARIGEON.

## Abstract

To appreciate the function of an organ, it is often critical to understand the role of rare cell populations. Unfortunately, this rarity often makes it difficult to obtain material for study. This is the case for the Drosophila male accessory gland, the functional homolog of mammalian prostate and seminal vesicle. In Drosophila, this gland is made up of two morphologically distinct cell types: the polygonally-shaped main cells, which compose 96% of the organ, and the larger, vacuole-containing secondary cells (SCs), which represent the remaining 4% of cells (~40 cells per lobe). Both cell types are known to produce accessory gland proteins (Acps), which are important components of the seminal fluid and are responsible for triggering multiple physiological and behavioral processes in females, collectively called the post-mating response (PMR). While a few genes are known to be specific to the SCs, the relative rarity of SCs has hindered the study of their whole transcriptome. Here, a method allowing for the isolation of SCs is presented, enabling the extraction and sequencing of RNAs from this rare cell population. The protocol consists of dissection, protease digestion and mechanical dissociation of the glands to obtain individual cells. Then, the cells are sorted by FACS, and living GFP-expressing SC singulets are isolated for RNA extraction. This procedure is able to provide SC-specific RNAs from ~40 males per condition in the course of one day. Given the speed and low number of flies required, this method enables the use of downstream RT-qPCR and/or RNA sequencing to the study gene expression in the SCs from different genetic backgrounds, ages, mating statuses or environmental conditions.

**SUMMARY:** Here, we describe the dissociation and sorting of a specific cell population from the Drosophila male accessory glands (Secondary cells), followed by RNA extraction for sequencing and RT-qPCR. The dissociation consists of dissection, proteases digestion and mechanical dispersion, followed by FACS purification of GFP-expressing cells.

## INTRODUCTION

Organs are composed of multiple cell types that have discrete functions and express different sets of genes. To get a precise idea of the functioning of an organ as a whole, it is critical to study the distinct cell types that compose this organ. Transcriptome analysis is a powerful approach to tackle cell function, providing a snapshot of genome expression and revealing active processes and pathways. But accessing the transcriptome of rare cell populations without contamination from more-abundant neighboring cells can be challenging. *Drosophila* accessory glands are a simple, secretory organ made up of only two secretory cell types. The rarer of the two cell types accounts for only 4% of the cells of this gland. For this reason, it has been difficult to access the full transcriptome and function of these cells.

Accessory glands (AGs) are key components of the male reproductive tract in insects, being responsible for the production of most of the seminal fluid proteins (SFPs) that are known to induce the physiological and behavioral processes in females collectively called the post-mating response (PMR). The PMR includes, but is not restricted to, increased ovulation and egg-laying, sperm storage and release, diet changes and gut growth, and decreased receptivity to secondary courting males ^1,2^. As such, AGs and SFPs are topics of intense interest to better understand the basic biological questions related to mating, reproduction, and evolution. Also, they have important impacts on major societal issues related to human health (some insects are vectors of deadly diseases) and agriculture (insects are both considered pests and critical for pollination and soil quality). *Drosophila melanogaster* is a prominent model for the study of AGs and ACPs, having brought many insights into the biology of these organs and the role of individual proteins regarding the PMR. The discoveries in fruit flies have largely affected the work in other species such as the disease vector *Aedes aegypti* ^3,4^, and other insects ^1,5^. Furthermore, the fact that the AGs secrete SFPs to be transferred to females during mating ^1,6^ makes the AGs the functional analog of mammalian prostate gland and seminal vesicle. Due to the functional and molecular similarities between the two tissue-tpyes, the AGs have been used as a model for the prostate gland in flies ^7^.

*Drosophila* accessory glands are composed of two lobes consisting of a monolayer of secretory cells surrounding a central lumen, and wrapped by smooth muscles. The secretory cells comprise two morphologically, developmentally and functionally distinct cell types: most of the gland is composed of polygonally-shaped main cells (~96% of the cells), while larger and rounded secondary cells (SC), make up the remaining 4% (~40 cells per lobe). Both cell types produce distinct sets of ACPs and work interdependently to induce and maintain the PMR.

The major trigger of the PMR is the Sex Peptide, a small 36 amino acid protein secreted by main cells and known to cause most PMRs in females ^8–10^. But many other ACPs produced by main and Secondary cells also affect different aspects of the PMR ^11–17^. Based on our current knowledge, it seems that SCs and their products are required to perpetuate the effects of SP past one day ^18^.

Thus far, most of the knowledge that we have accumulated on SC biology comes from candidate approaches, finding the expression of one particular gene or protein in these cells and determining its role in the development and/or function of the SCs. These genes include the homeodomain protein *Defective proventriculus* (*Dve*, ^19^), the lncRNA *MSA* ^20^, *Rab6, 7, 11* and *19 ^21^*, *CG1656* and *CG17575* ^11,15,21^ and the homeobox transcription factor *Abdominal-B* (*Abd-B*) ^18^. A mutant line deficient for both the expression of *Abd-B* and the lncRNA *MSA* in secondary cells has been used to determine that secondary cells are required for Sex Peptide to be properly stored in the female reproductive tract, which results in a shortened PMR (from ~10 days to one day) ^12,18,20^. At the cellular level, the SC of this mutant almost lose their characteristic vacuole like structures ^18,20,21^. This mutant line was successfully used to identify some genes involved in these phenotypes by comparing transcriptomes of wild type versus mutant accessory glands ^12^.

Unfortunately, it was difficult to access the full genetic program of SC, because of their relative rareness in an organ made up primarily of main cells. For qPCR validation of genes suspected to be induced under particular conditions, the abundance of main cells would often hide the variation in gene expression, and performing *in situ* hybridization on glands proved to be tricky. We thus decided to develop a method for isolating purified SC RNA that was easy and fast enough to perform in a variety of different genetic backgrounds or environmental conditions.

*Abd-B* and *MSA* expression in SCs relies on the *D1* enhancer, a 1.1kb piece of DNA located in the iab-6 regulatory region of the Bithorax complex ^18,20^. GAL4 drivers containing this sequence are expressed in secondary cells and, when associated with a UAS-GFP, give a strong GFP signal in live SCs, allowing clear visualization and FACS sorting of these cells. The *iab-6*^*cocuD1*^ chromosome has a small deletion of this specific *D1* enhancer, abrogating *Abd-B* and *MSA* expression in SC, and causing the phenotypes described above ^20^. We performed this protocol on *wt* and *iab-6*^*cocuD1*^ mutant accessory glands as a proof of principle that this approach can not only provide the wild type transcriptome of this rare cell type, but also to identify mis-regulated genes involved in SC function.

## PROTOCOL

### 1. Drosophila line generation and male collections

1.1. Sorting SCs using this protocol requires that males express GFP in the SCs but not in main cells. Use an AbdB-GAL4 or other appropriate driver (described in ^18^) recombined with UAS-GFP (on chromosome 2) for this method. Other mutations can be added along with these transgenes, if required.
1.2. Collect healthy, virgin males and place them into vials with food in groups of 20-25. Typically, the protocol requires 2 batches of 20 males for each genotype. NOTE: The age, mating status, food and social environment affect accessory gland biology. To control for these parameters, males aged for 3-4 days after pupal eclosion at 25°C with a 12h/12h light/dark cycle, on regular food with yeast, in groups of 20-25 males were used here.

### 2. Solutions and material preparation

#### 2.1. Solutions and aliquots

2.1.1. Prepare Serum Supplemented Medium (SSM). Add 10% heat inactivated fetal bovine serum and 1% Penicillin-Streptomycin into Schneider’s Drosophila medium.
2.1.2. Prepare aliquots of 1X TrypLE Express Enzyme, keep at room temperature.
2.1.3. Prepare aliquots of papain [50U/mL], store at −20 °C and thaw only once.
2.1.4. Prepare 1X PBS, keep at room temperature.

#### 2.2. Material

2.2.1. Standard lab equipment including micropipettes, tips, Eppendorf tubes, 24-well plates, single depression and 3 depressions “glass spot plates”, ice buckets and Bunsen burners should be accessible.
2.2.2. Thermo shaker (37°C 1000rpm, then 65°C).
2.2.3. Fine dissection forceps.
2.2.4. Binocular for accessory gland dissection.
2.2.5. Binocular with fluorescence source to see GFP.
2.2.6. Low retention pipet tips (200μl and 1000μl).
2.2.7. Fluorescence Activated Cell Sorter.

#### 2.3. Flame-round multiple pipet tips for handling and physically dissociating accessory glands

NOTE: Using low binding tips is important for handling accessory glands because they tend to adhere to untreated plastic. Flame rounding narrows the tip opening while smoothening the edge of the tip. This is critical to ensure smooth yet efficient dissociation after peptidase digestion.

2.3.1. Cut one 200μl tip with a razorblade and flame round it so that the opening is wider and smooth for handling at step 3.7.
2.3.2. For mechanical trituration (steps 5.2. and 5.3.) the opening of 1000μl tips is reduced. Prepare a few 1000μl tips by putting the narrow opening near the Bunsen flame for less than one second. Rotating helps avoiding clogging or over-melting. Test them by pipetting water and sort them from wider to narrower based on the speed of aspiration. Test the efficacy of each tip by using them on a small scale sample before using them for the real experiment (especially the narrowest tips). Verify that the tip allows complete dissociation of a treated sample while preserving cell viability (see below). NOTE: Good flame-rounded pipet tips are washed with water at the end of the procedure, dried and reused to provide reproducible dissociation, from one day to another.

### 3. Accessory glands dissection

3.1. Place 20-25 males in a glass dish on ice.
3.2. Dissect one male at a time in SSM. Reproductive tracts are taken off, and AG pairs are cleared from other tissues with the exception of the ejaculatory duct. NOTE: The testes should be removed, as the released sperm can create clumps that disturb the dissociation process. The ejaculatory bulb must also be removed, as it tends to float, making handling more difficult.
3.3. Using forceps, transfer AGs to a glass “3 spot plate” containing SSM (room temperature). Repeat steps 3.2 and 3.3. until 20 pairs of AGs are pooled in a single well with SSM. NOTE: This step should be achieved in 15-20 minutes. Glands should look healthy, and the muscle layer around glands should continue their peristaltic movements. GFP can be checked with a fluorescent binocular. NOTE: Dissecting batches of 20 males is a good balance between manageability and efficiency; flies do not spend too much time on ice and the glands do not spend too much time in SSM prior dissociation. Working with smaller sample of 5 to 10 males is good for troubleshooting, but results in more cell loss during transfers (from plate to tube to plate to dissociation to tube).
3.4. Transfer AGs into 1X PBS at room temperature for 1-2 minutes. NOTE: Rinsing glands with PBS results in better dissociation. However, this step should be short as accessory glands seem to be stressed in this solution. Placing dissociated secondary cells into PBS results in a rapid increase in SC size followed by the death of most SC, indicating that PBS is not an homeostatic solution for these cells.
3.5. Prepare dissociation solution by diluting 20μl papain [50U/mL] into 180μl TrypLE 1X (i.e. 9μl TrypLE + 1μl papain per male). Transfer AGs into this solution.
3.6. Isolate the distal part of AGs (containing SCs) from the rest by pinching and cutting with fine forceps. Hold the middle of the gland firmly and cut with one sharp tip of the other forceps. Remove ejaculatory duct and the proximal part of the glands to improve dissociation and reduce cell sorting time.
3.7. When all gland tips are isolated, carefully transfer them to a 1.5 mL Eppendorf tube using a pre wet cut and flame-rounded low binding 200μl tip (cf. 2.3.1) NOTE: Dissection of 20 pairs of gland tips should take ~15-20 minutes, during which digestion by Trypsin and Papain starts at at room temperature. NOTE: Always pre-wet tips with the appropriate solution, and rinse them carefully between each sample by pipetting up and down twice to avoid contamination. Keep a dedicated tube of TrypLE and a tube of SSM for this purpose. NOTE: One single trained person can process 4×20 males in one morning, to obtain secondary cells’ RNAs from 2 conditions in one day. However, two (or more) persons can participate in the initial steps of accessory glands collection (3.1. to 3.3.) in order to obtain several samples from the same day, which is desired for comparing multiple samples. Preferentially, steps 3.4. onwards should be performed by a single experimenter to improve reproducibility.

### 4. Cells dissociation

4.1. Place the tube in a Thermo Shaker at 37°C with 1000rpm rocking, for 60 minutes. NOTE: The digestion time and the agitation are both critical for the success of the dissociation. Reducing time or immobile digestion will result in poor dissociation, probably because accessory gland cells are protected from peptidases by the outer muscle layer and inner viscous seminal fluid.
4.2. Stop the digestion by adding 1mL of SSM (room temperature). Quickly proceed to step

### 5. Mechanical disruption

5.1. Transfer each sample to a well in a 24-well plate using a wide, rounded 1000μl tip, pre-wet with SSM. Check GFP fluorescence under the binocular, some SC should be detached but most gland tips should look intact.
5.2. Mechanically disrupt the gland tips by pipetting up and down 3-5 times with a rounded narrow pipet tip.
5.3. Repeat the process of pipetting up and down 1-2 times with a rounded very narrow pipet tip. NOTE: Check fluorescence and repeat the mechanical disruption process if required. After step 5.2. big tissue patches should be broken up. After step 5.3., individual cells should predominate. Once a few samples have been processed and satisfactory result have been achieved (perfect-looking dissociation with healthy looking secondary cells), these pipetting steps can be performed in the Eppendorf tubes for practical reasons. The use of 24-well plates is good as a first step because they allow for an easy monitoring of the process.
5.4. Let the cells settle down in the plastic well for ≥15 minutes, and remove the excess SSM to reduce the FACS time. NOTE: Letting cells settle proved superior to alternative approaches such as centrifugation, and allows one to visually inspect the cells to make sure they are not lost in the process.
5.5. Pool the identical samples (2 batches from 20 males each) and transfer them into a clean 1.5mL tube. Rinse the well with a small volume of SSM to retrieve remaining cells, and add them to the tube.
5.6. Prepare tubes containing 300μl Cell Lysis Solution with 1μl [50μg/μl] Proteinase K for each sample (provided in the RNA extraction kit). Prepare one tube for the main cells and one for the secondary cells sorted from each sample.

### 6. Fluorescence-activated Cell Sorting (FACS)

6.1. Add 0.3mM Draq7 (viability marker) to each sample to be sorted. This can be done a few minutes ahead of time. CAUTION: Draq7 should be handled with caution.
6.2. Sort GFP-positive Draq7-negative cells. They should be a homogeneous population of live Secondary cells. In a different tube, sort a homogeneous population of smaller GFP-negative Draq7-negative cells. These should be the main cells. We used the following gating strategy for FACS:
  6.2.1. Select total cells based on FSC-SSC in order to exclude debris.
  6.2.2. Remove dead cells by gating out Draq7-positive cells. Draq7 was excited with a 640 nm laser and fluorescence emission was collected with a 795/70 band-pass filter.
  6.2.3. Exclude doublets using a double gating on FSC-A vs FSC-H and SSC-H vs SSC-W.
  6.2.4. Sort ~550 GFP-positive cells into a 1.6 mL Eppendorf tube containing Lysis buffer with Proteinase K, they are considered Secondary cells (SC). GFP was excited at 488 nm and fluorescence emission was collected with a 526/52 band-pass filter.
  6.2.5. Sort ~1000 non-GFP-positive cells composing a population of small cells homogeneous in size into a 1.6 mL Eppendorf tube containing Lysis buffer with Proteinase K, they are considered main cells (MC).
  6.2.6. Vortex samples.

NOTE: The pressure on the MoFlo Astrios was set at 25 PSI and the cells were passed through a 100µm nozzle. The rate of sorting was about 2.000cells/sec.

NOTE: Stringently remove cell doublets using a double gating, as main cell contamination must be reduced as much as possible.

NOTE: From 40 males (~40*80 = 3200 SC), yields generally range from 600 to 800 live, singulet secondary cells (around 20-25% efficiency). We stop sorting around 550 Secondary cells and 1000 Main cells to normalize samples.

NOTE: The cells can be sorted into RNAlater instead of lysis buffer if one wants to extract RNAs another day of pool samples from different days. We did not do this ourselves.

### 7. RNA extraction

NOTE: RNA extraction is performed using Epicentre MasterPure RNA Purification Kit, with following adaptations. Other kits might be used as long as the yield is high enough to prepare a library for sequencing from ~500 cells (2ng RNAs in our case).

7.1. Cell lysis: cells in lysis buffer containing proteinase K are heated at 65°C for 15 minutes. Vortex the samples every 5 minutes.
7.2. Place samples on ice for 5 minutes, and proceed with nucleic acid precipitation, following manufacturer’s recommendations in: “Precipitation of Total Nucleic Acids” and “Removal of Contaminating DNA from Total Nucleic Acid Preparations”.
7.3. Resuspend the RNA pellet in 10μl RNase-free TE buffer.
7.4. Add 1μl RNases inhibitor (optional).
7.5. Keep samples at −80°C.

### 8. Quality controls (RNA quality, quantity, specificity)

8.1. Estimate RNA quality and concentration. Due to the small volume and concentration, we use Agilent RNA 6000 Pico chips. Good quality RNA is defined as non-degraded, visible as a low baseline with sharp peaks corresponding to rRNAs.
8.2. RT-qPCR to control the identity of sorted cells
  8.2.1. Reverse transcription is performed using 2ng total RNAs using random hexamers as primers. Perform RT on Secondary Cell RNA and Main Cell RNA. NOTE: cDNAs can be diluted, aliquoted, and kept frozen for later use.
  8.2.2. Real Time quantitative PCR is performed using appropriate primer pairs to quantify housekeeping genes (alpha-Tubulin, 18S rRNA), Secondary cell specific genes (MSA, rab19, Abd-B) and Main cell specific gene (Sex peptide).

### 9. Sequencing (Library preparation, Sequencing, data analysis)

9.1 cDNAs are synthesized from 2ng total RNAs with polydT primers, using the SMARTer technology allowing subsequent amplification (https://www.takarabio.com/products/cdna-synthesis/cdna-synthesis-kits/smarter-cdna-synthesis-kits).
9.2 Preparation library using Nextera XT kit (https://emea.illumina.com/products/by-type/sequencing-kits/library-prep-kits/nextera-xt-dna.html).
9.3 Sequence with multiplexed, single reads of 100 nucleotides (although 50 nucleotides reads are enough for most purposes). We obtained around 30 million reads and a yield 2500–3000 Megabases per sample.

### 10. Data analysis

10.1. FastQC (http://www.bioinformatics.babraham.ac.uk/projects/fastqc/).
10.2. Reads are mapped to the reference Drosophila genome (UCSC dm6) using the STAR aligner and .bam files are generated for subsequent visualization of reads on Integrative Genomics Viewer (IGV).
10.3. Gene count was performed using HTSeq.
10.4. Normalization, statistical analysis of differential expression, PCA and MA plots were performed using edgeR. The Trimmed Mean of M-values (TMM) method was used to normalize gene counts ^22^.
10.5. Statistical analysis of gene expression using General Linear Model, quasi-likelihood F-test with False Detection Rate (FDR) and Benjamini & Hochberg correction (BH).

## REPRESENTATIVE RESULTS

The method presented here allows, in the course of one day, for isolating GFP-expressing secondary cells from Drosophila accessory glands, and extracting their RNA for sequencing.

**Figure 1:**
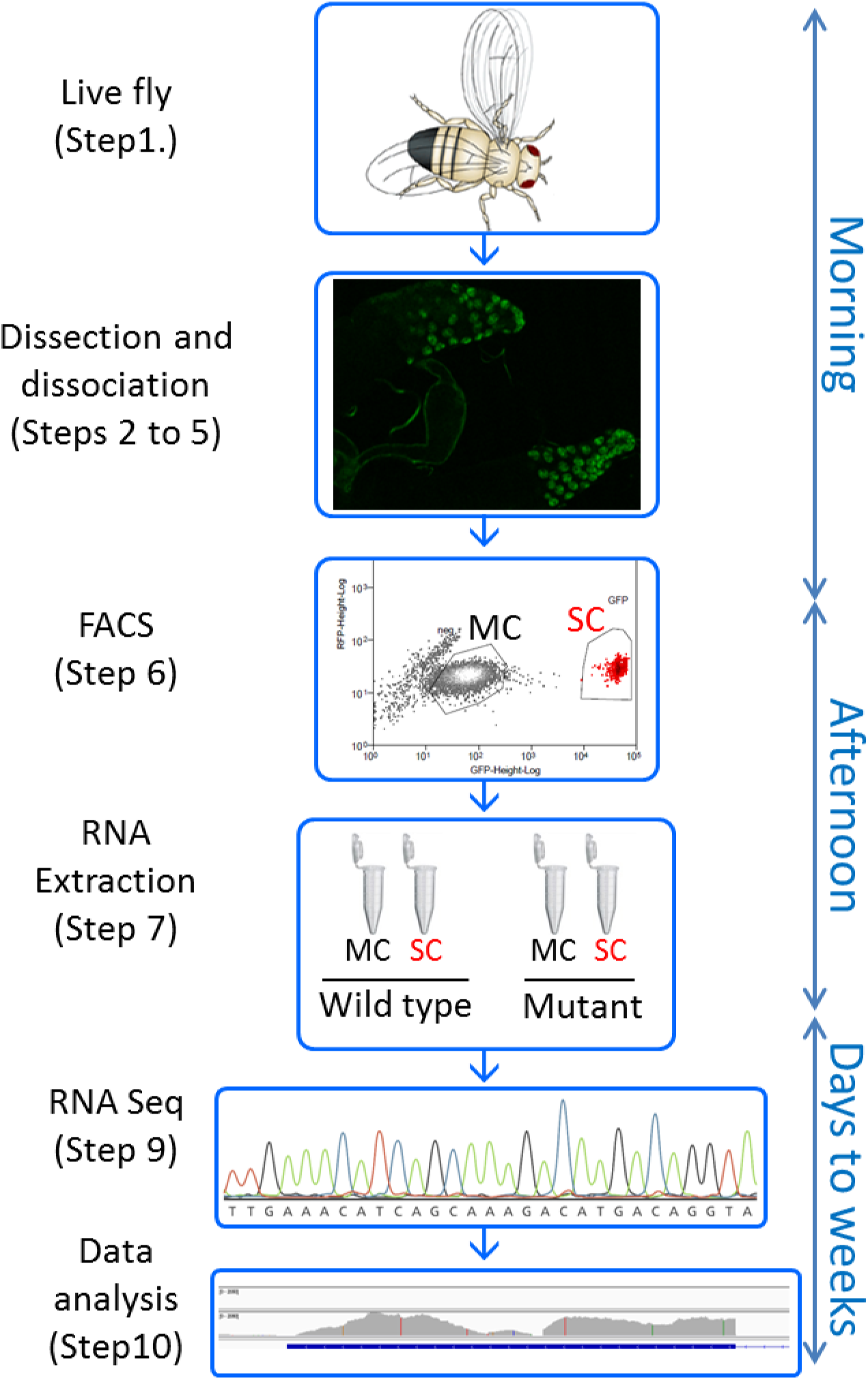
Overview of the protocol. Key steps of the protocol are shown, with the timeline on the right side. This procedure allows one starting with live Drosophila in the morning to have dissociated accessory gland cells by noon, sort them based on GFP expression, and get their RNAs extracted by the end of the working day. RNA sequencing and data analysis will typically take a few weeks.

We use here the *Abd-B-GAL4* construct described in ^18^ to express GFP in secondary cells (SC) but not in main cells (MC) (Figure 2A). The first objective of the method is to get the transcriptome of “wild type” SC (wild type is put in quotation marks because these flies are transgenic animals carrying a GAL4 driver express GFP). The second objective is to be able to obtain SC RNA quickly and easily enough so it is possible to compare their transcriptomes in different conditions. To test for this, we performed this protocol from “wild type” and “*iab-6*^*cocuD1*^” mutants carrying a deletion of 1.1kb removing the SC enhancer of *Abd-B* as well as the promoter of the *MSA* transcript, known to be critical for SC development, morphology and function ^18,20^ (Figure 2B and C). We repeated this protocol 3 times on different days to generate the triplicates of each genotype presented here throughout the figures (hereafter referred as wt-1,-2,-3 for the *wild type* and D1-1,-2 and −3 for the *iab-6*^*cocuD1*^).

**Figure 2:**
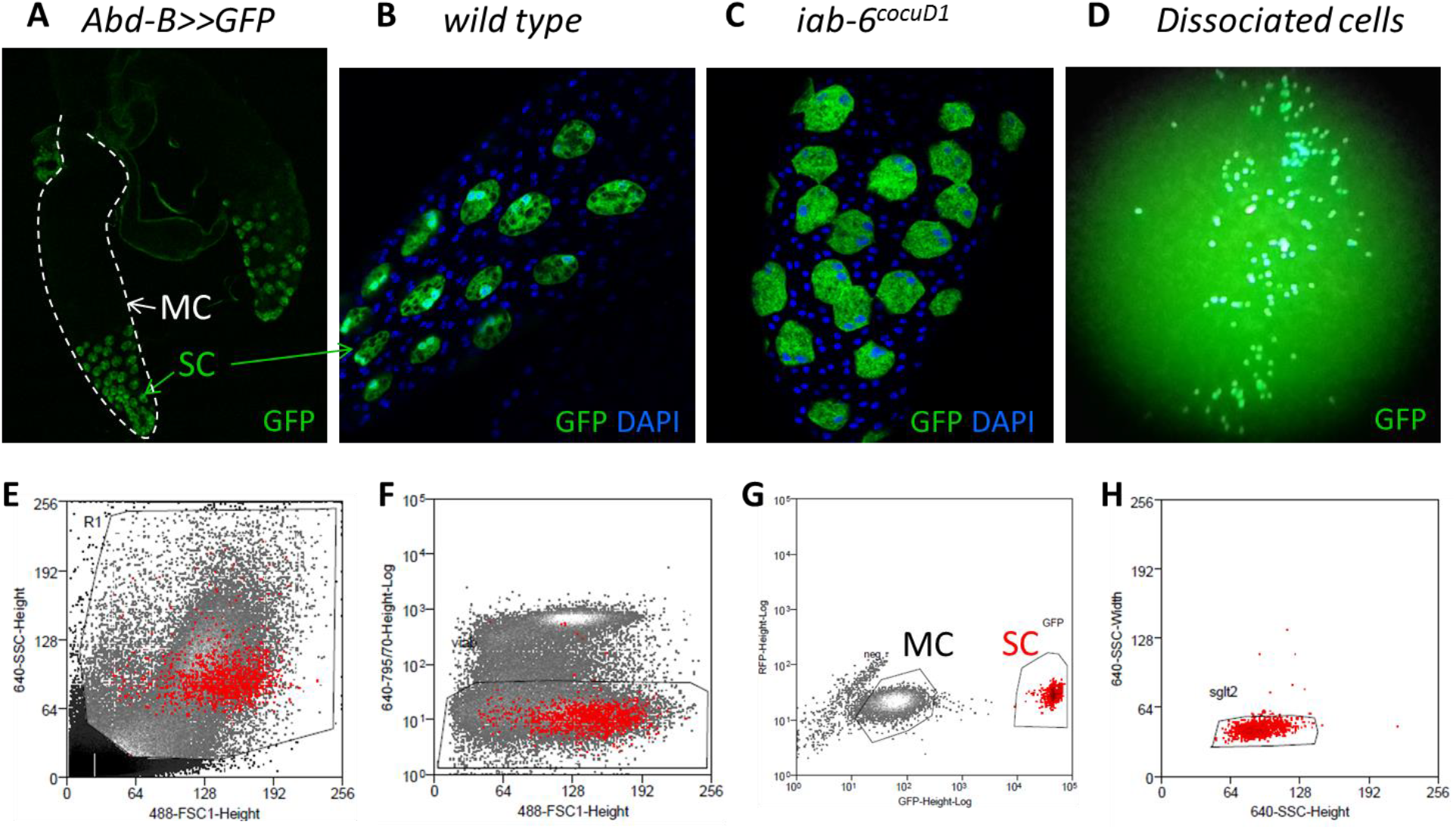
Isolating and sorting GFP-expressing secondary cells. A-Confocal image of *Abd-B:GAL4 UAS-GFP* accessory gland expressing GFP in secondary cells (SC), but not in main cells (MC). B and C show enlarged views of accessory gland distal part with SC expressing GFP, in wild type (B) or *iab-6*^*cocuD1*^ (C) background. Panel D shows dissociated cells at low magnification under the GFP binocular. Panels E to H displays FACS gating strategy to purify SC and MC. Red dots on all panels show SCs as defined by GFP expression (G). First, debris are excluded (E, step 6.2.1.) as well as the Draq-7 positive dead cells (F, step 6.2.2.). GFP-positive cells are selected (SC) as well as an homogeneous population of GFP-negative cells (MC) (G, steps 6.2.4. and 6.2.5.). Doublets are excluded from both MC and SC population (step 6.2.3., only the SSC-H vs SSC-W gating for SC is shown on panel H as an example).

The method described herein allows dissociating MC and SC from *Drosophila* accessory glands in only a few hours (Figure 2D). These cells are then sorted by FACS into two distinct tubes to isolate MCs and SCs. The FACS gating strategy is presented in Figure 2E to G. The addition of Draq7 allows the estimation cell viability, which is around 70% for the whole sample (Figure 2F). The singulets - estimated by double gating on FSC-A vs FSC-H and SSC-H vs SSC-W - represent over 90% of the SC population, and over 80% of the MC population, reflecting the efficiency of dissociation (exemplified in Figure 2H). 10 to 12% of SC sortings were aborted due to the presence of another cell or debris in the droplet. Starting with 40 males, we typically stop sorting after collecting 550 SCs and 1000 MCs. With one out of 6 samples, we did not reach the objective of 550 SC: the wt-1 sample was prepared with only 427 SC, but gave equally good results.

SC and MC are sorted into lysis buffer and RNA extraction is performed as soon as all samples are ready in order to obtain RNA pellets by the end of the same day. RNA quality and quantity are estimated on a Bioanalyser using an appropriate chip to work with small volumes and low concentrations. Since estimated concentrations were quite variable between samples (ranging from 344pg/μl to over 1300pg/μl, see Figure 3A), the starting material for both RT-qPCR and cDNA library synthesis was roughly adjusted to 2ng (the measured concentration is not very precise). The expression of specific genes was quantified by Real Time qPCR on SC and MC extracts of the “wt” condition to control for the identity of the sorted populations. The results shown in Figure 3B are relative quantifications normalized to *alpha-tubulin* for each sample. As expected, housekeeping genes *alpha-tubulin* and *18S* rRNA are detected in all samples. On the contrary, the SC-specific gene *rab19* is only detected in SC extracts, while the MC-specific gene Sex Peptide is only detected from MC. As expected, the SC-specific transcript *MSA* whose promoter is deleted in *Iab-6*^*cocuD1*^ is detected only from wt SC and not from mutant SCs. Together, the quality controls presented in Figure 3 show that the RNAs obtained with this procedure are not degraded and that SC and MC populations are successfully sorted, from both wt and mutant accessory glands. We sequenced only secondary cells’ RNAs, but note that this protocol allows for concomitant RNA profiling of both SC and MC.

**Figure 3:**
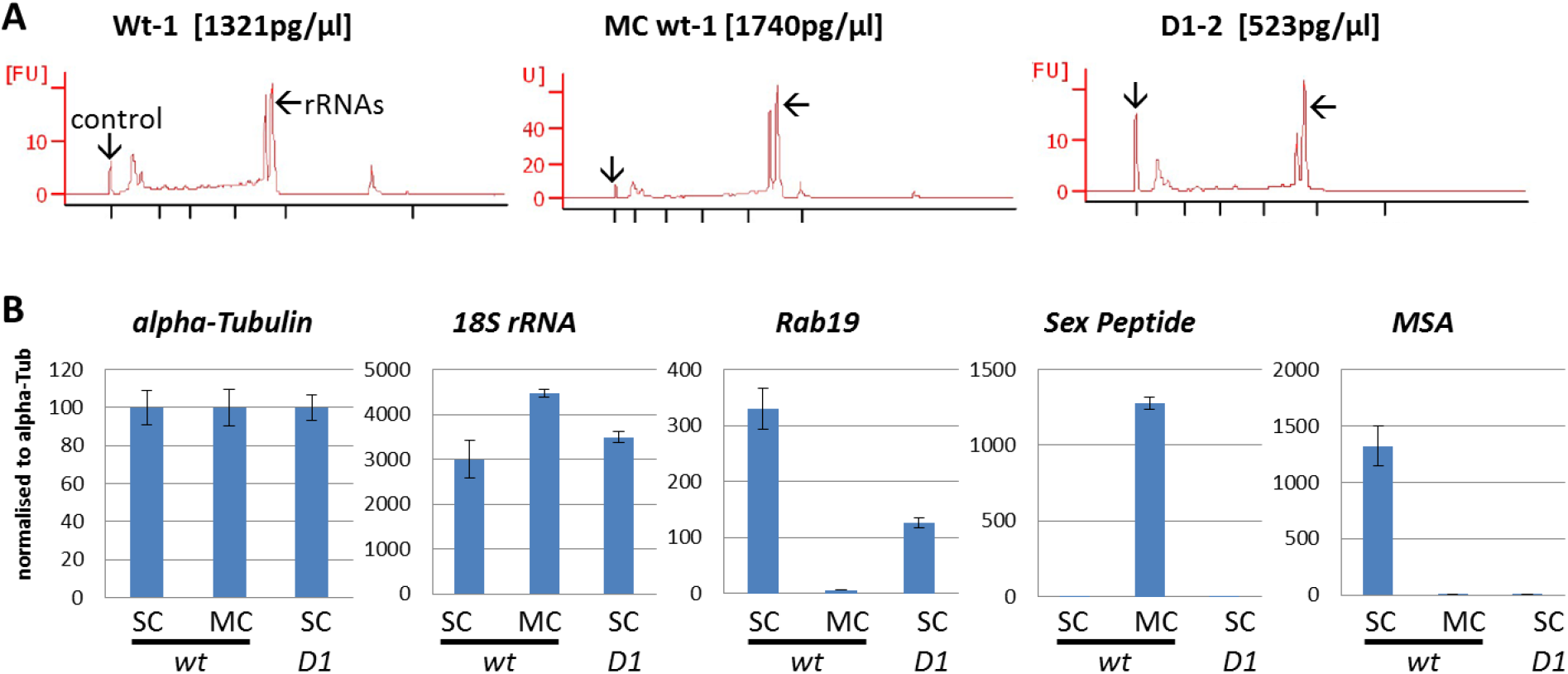
QC on RNAs: quality, quantity, and cell type specificity. A- Control of RNA quality and concentration estimation on PicoChip. The 25nt peak is the control for quantification. Low baseline indicates low degradation and the 2 major peaks are the ribosomal RNAs. The 3 samples used for RT-qPCR are shown, their quality is representative of all RNA samples used in this study, and their estimated concentrations reflect the variation in total RNA we obtained between samples. B- RT-qPCR demonstrate the specificity of the sorting of secondary cells (SC) and main cells (MC). Gene expression quantification is done using the q=2^(40-Cq) formula. Each triplicate of each gene in each condition is normalized to the mean quantity of alpha-Tubulin RNA to compensate for total RNA variation. Error bars show standard deviation. Wt means wild type and D1 refers to the iab-6^cocuD1^ mutant.

The RNA-sequencing was performed using standard protocols. For the purpose of this method, we will only discuss here the pertinent quality control analysis. After sequences were obtained, the reads were mapped to the reference genome, attributed to genes, counted and normalized. Principal Component Analysis (PCA) was performed on all 6 samples (three “*wild type*” replicates and three *iab-6*^*cocuD1*^ replicates). PC1 accounts for as much of the variability in the data as possible, and PC2 accounting for as much of the remaining variability as possible. As presented in Figure 4A, the 3 *wt* replicates cluster together, and far away from *Iab-6*^*cocuD1*^ samples, showing that *wt* samples are similar to each other, but different from mutant samples. This shows the reproducibility of the method, and its ability to characterize the divergent genetic program of mutant SC. We note that while D1-2 and D1-3 samples cluster together, the D1-1 sample is quite different. As all quality controls for this sample are good and similar to all other samples, we can exclude a problem in sample preparation (29 million total reads of which >76% align uniquely to the genome and >77% of them are attributed to a gene, >90% mRNAs, <3% rRNA). This variance could thus reflect that gene expression in *iab-6*^*cocuD1*^ SC is unstable, although more replicates would be necessary to test this hypothesis.

**Figure 4:**
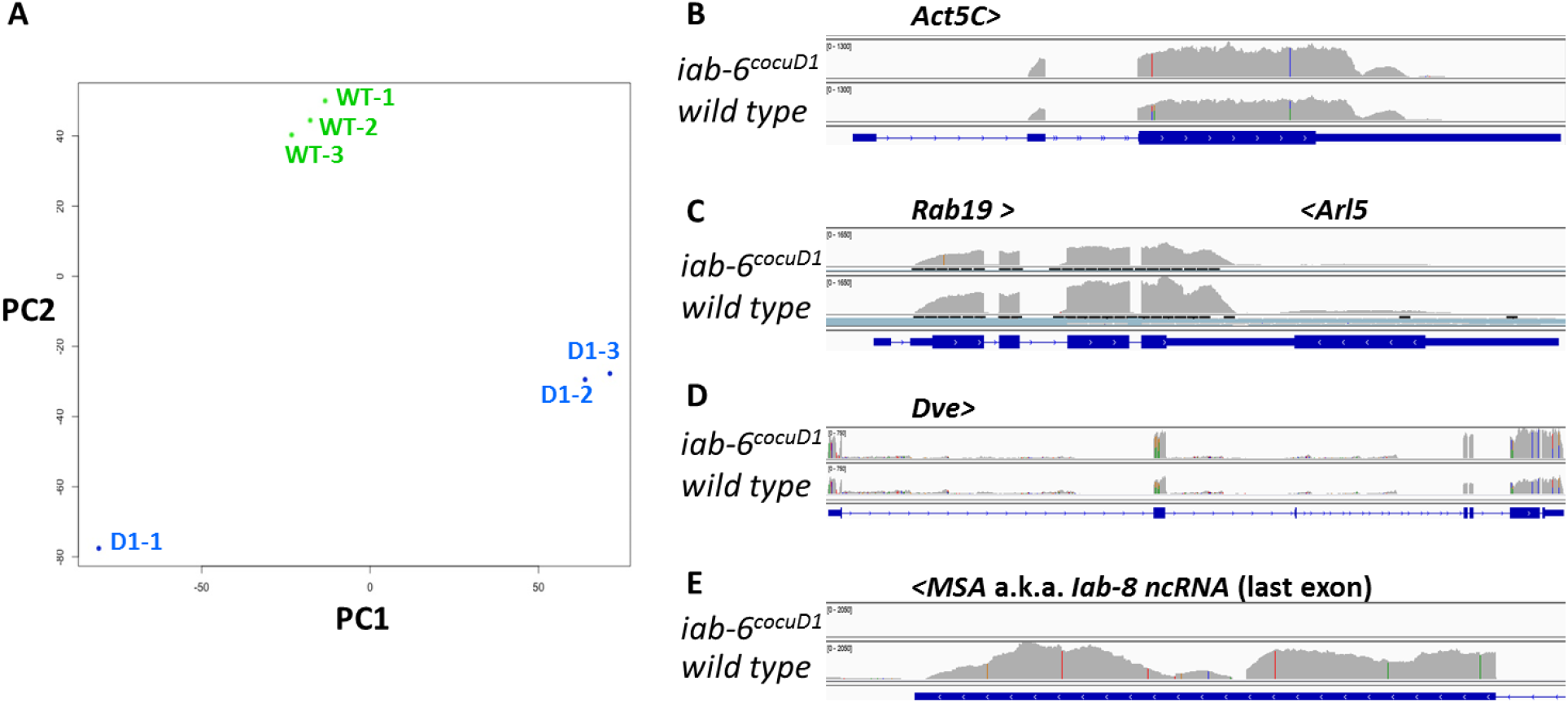
QC on RNA sequencing data. A-Principal Components Analysis (PCA) on wt-1,-2,-3 (green dots) and D1-1,-2,-3 (blue dots) RNA sequencing datasets. Panels B to E show sequencing reads mapped to the Drosophila reference genome using the IGV software. Only one representative sample of each genotype is shown for clarity sake (wt-1 and D1-2), and only a few specific loci are shown. Gene names are written on top of each panel, > and < symbols refer to their orientation. Numbers in brackets represent for each track the scale for the number of reads per DNA base pair. This scale is the same for both conditions for a given gene, but varies between genes for better visualization. Blue bars at the bottom of each panel show genes’ introns (thin line), exons (wide line) and ORF (rectangles with >>). Note that Rab19 and Arl5 are overlapping, convergent genes (C).

Visualization of reads on the genome at particular genes allows a direct and visual estimation of the quality of the data. In Figure 4 a selection of genes is shown, with one representative replicate of each genotype. As expected housekeeping genes such as *Act5C* (Figure 4B) are expressed in both conditions, as well as the SC genes *Rab19* and *Dve* (Figure 4C-D). The absence of reads in introns shows that polydT reverse transcription successfully selected the mature spliced mRNAs to prepare cDNA library. Importantly, we can see strong and significant variations in some genes’ expression levels comparing *wild type* and *iab-6*^*cocuD1*^. This is exemplified by the *MSA* gene presented in Figure 4E whose expression is strong in *wild type* and absent from *iab-6*^*cocuD1*^. This gene is shown as a proof of principle that this method allows identifying significantly mis-regulated genes which could shed light on the mechanisms responsible for the phenotypes observed in this mutant and give new insights into normal SC function.

**Table 1:**
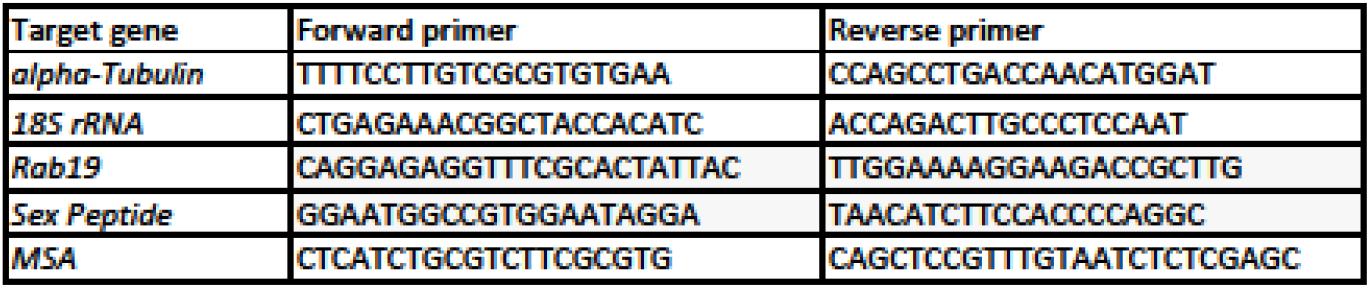
Primers sequence

**Table 2:** Materials used

| Name of Material/ Equipment | Company | Catalog Number | Comments/Description |
| --- | --- | --- | --- |
| 24-wells Tissue Culture plate | VWR | 734-2325 |  |
| 3 spots glass plate |  |  |  |
| Binocular for dissection |  |  |  |
| Binocular with light source for GFP |  |  |  |
| Bunsen |  |  |  |
| Draq7 0.3 mM | BioStatus | DR71000 |  |
| eppendorf tube 1.6 mL |  |  |  |
| FACS | Beckman Coulter | MoFlo Astrios |  |
| Fine dissection forceps |  |  |  |
| foetal bovine serum | Gibco | 10270-106 | heat inactivated prior to use |
| Glass dish |  |  |  |
| ice bucket |  |  |  |
| ImProm-II Reverse Transcription System | Promega | A3800 |  |
| MasterPure RNA Purification Kit | Epicentre | MCR85102 |  |
| Nextera XT kit | illumina |  |  |
| P1000 | Gilson |  |  |
| P20 | Gilson |  |  |
| P200 | Gilson |  |  |
| Papain |  |  | 50U/mL stock |
| PBS |  |  | home made |
| Penicillin-Streptomycin | Gibco | 15070-063 |  |
| polydT primers |  |  |  |
| RNA 6000 Pico kit | Agilent |  |  |
| Schneider's Drosophila medium | Gibco | 21720-001 |  |
| smarter cDNA synthesis kit | Takara |  |  |
| SYBR select Master mix for CFX | applied biosystems | 4472942 |  |
| Thermo Shaker | Hangzhou Allsheng intruments | MS-100 |  |
| Tipone 1250 µl graduated tip | Starlab | S1161-1820 |  |
| Tipone 200 µl bevelled tip | Starlab | S1161-1800 |  |
| TrypLE Express Enzyme | Gibco | 12604013 |  |
| Vortex |  |  |  |

## DISCUSSION

Secondary cells represent a minor cell type of the *Drosophila* accessory gland, yet play a critical role in male fertility by maintaining the post-mating response in females. In this protocol, we present a quick and efficient method to access the full transcriptome of these cells that account for only 4% of the organ, i.e. ~80 cells per individual. This method is based on dissection, peptidase digestion and FACS sorting, and can be performed in one day for multiple samples (except the RNA sequencing part).

Methods to dissociate cells from *Drosophila* tissue such as imaginal discs were described previously ^23^. However, our attempts to simply use these methods with accessory glands failed, prompting us to develop this method. For a successful dissociation, the peptidases have to access the accessory gland cells, which are protected by an outer muscle layer and an inner viscous seminal fluid. Thus, cutting the glands open (step 3.6.) and digesting for at least 60 minutes with vigorous shaking (step 4.1.) is critical to success. TrypLE allowed gentle dissociation, conserving the integrity of the gland and viability of the cells until mechanical dissociation. While papain and collagenase were not sufficient to dissociate the cells, we found that both of those enzymes improved the dissociation in association with TrypLE (less pipetting was required to obtain perfect dissociation, resulting in better survival). The trituration with narrow, rounded tips (steps 5.2. and 5.3.) is a key step and should be optimized in pilot experiments due to the way these tips are created (see step 2.3.2. and the Note after step 5.3. for guidelines).

This method allows one to isolate around 500-800 individual, live Secondary cells from 40 males (we stop sorting around 550 cells per sample to normalize material for RNA extraction). This corresponds to ~20% (±5%) efficiency assuming a starting material containing ~ 3200 SC (40 males × 80 SC). 20% recovery was enough for our purpose as we could obtain several samples in one day. However it might be improved by different means, including: working in larger batches and reducing transfers (doing trituration in the digestion tube, skipping step 5.4. and going straight to the FACS); performing a more gentle trituration (a significant proportion of dissociated GFP+ cells visible after step 5.3. die in the hour following trituration, as they are probably damaged in the process); reducing the digestion duration (using a more concentrated TrypLE might be an attractive option); reducing the stringency of singulet selection (step 6.2.3.); reducing time between time between dissociation and FACS…

Secondary cells have unique cell morphology, having two polyploid nuclei and a vacuole-filled cytoplasm, consistent with a role in producing, modifying, and secreting a large quantity of proteins into the seminal fluid ^24^. These secretory cells fulfill non-redundant functions essential for male fecundity ^12,20,25^, and having a global view of all the genes that they express gives new insights into their normal function. We show here that the transcriptome of these cells can be obtained from a relatively small number of males, allowing for the comparison of different conditions. Here we use one mutant known to affect secondary cells morphology and function, and show that its transcriptome is significantly changed, suggesting that this method will allow identification of new important secondary cell genes. Previously in the lab a similar analysis of *wt* and *iab-6*^*cocuD1*^ SC had been done by manually picking the cells, and produced good quality RNA sequencing data as well, but the method was too labor intensive to be extended to other conditions. In a separate manuscript, currently in preparation, we will extensively present and compare our datasets, and more importantly analyze them regarding their biological significance and how they help us understanding the normal function of SC.

As the morphology, vacuolar content and the number of Secondary cells has been shown to change with age, mating status, and diet ^21,26,27^, it would be interesting to compare SCs under different conditions. Having a simple and fast protocol will allow one to study SC under all of these different conditions. It is noteworthy that this protocol allows simultaneous isolation of main cells from the same individuals, and thus could also be used without modification to see how the different genetic and environmental parameters affect mains cells at the same time.

## ACKNOWLEDGMENTS

This work was funded by the State of Geneva (CI, RKM, FK), the Swiss National Fund for Research (www.snf.ch) (FK and RKM) and donations from the Claraz Foundation (FK).

We thanks the members of the Karch lab, Dr. Jean-Pierre Aubry-Lachainaye from the Flow Cytometry core facility of University of Geneva for setting up the FACS protocol, and the iGE3 genomics plateform for cDNA library preparation and RNA sequencing. We are grateful to Luca Stickley for his help to visualize the reads on IGV.

## REFERENCES

1 Avila, F. W., Sirot, L. K., LaFlamme, B. A., Rubinstein, C. D. & Wolfner, M. F. Insect seminal fluid proteins: identification and function. Annu Rev Entomol. 56 21–40, (2011).

2 Carmel, I., Tram, U. & Heifetz, Y. Mating induces developmental changes in the insect female reproductive tract. Curr Opin Insect Sci. 13 106–113, (2016).

3 League, G. P., Baxter, L. L., Wolfner, M. F. & Harrington, L. C. Male accessory gland molecules inhibit harmonic convergence in the mosquito Aedes aegypti. Curr Biol. 29 (6), R196–r197, (2019).

4 Degner, E. C. et al. Proteins, Transcripts, and Genetic Architecture of Seminal Fluid and Sperm in the Mosquito Aedes aegypti. Mol Cell Proteomics. 18 (Suppl 1), S6–s22, (2019).

5 Wedell, N. Female receptivity in butterflies and moths. J Exp Biol. 208 (Pt 18), 3433–3440, (2005).

6 Laflamme, B. A. & Wolfner, M. F. Identification and function of proteolysis regulators in seminal fluid. Mol Reprod Dev. 80 (2), 80–101, (2013).

7 Wilson, C., Leiblich, A., Goberdhan, D. C. & Hamdy, F. The Drosophila Accessory Gland as a Model for Prostate Cancer and Other Pathologies. Curr Top Dev Biol. 121 339–375, (2017).

8 Kubli, E. & Bopp, D. Sexual behavior: how Sex Peptide flips the postmating switch of female flies. Curr Biol. 22 (13), R520–522, (2012).

9 Kubli, E. Sex-peptides: seminal peptides of the Drosophila male. Cell Mol Life Sci. 60 (8), 1689–1704, (2003).

10 Liu, H. & Kubli, E. Sex-peptide is the molecular basis of the sperm effect in Drosophila melanogaster. Proc Natl Acad Sci U S A. 100 (17), 9929–9933, (2003).

11 Ram, K. R. & Wolfner, M. F. Sustained post-mating response in Drosophila melanogaster requires multiple seminal fluid proteins. PLoS Genet. 3 (12), e238, (2007).

12 Sitnik, J. L., Gligorov, D., Maeda, R. K., Karch, F. & Wolfner, M. F. The Female Post-Mating Response Requires Genes Expressed in the Secondary Cells of the Male Accessory Gland in Drosophila melanogaster. Genetics. 202 (3), 1029–1041, (2016).

13 Avila, F. W. & Wolfner, M. F. Acp36DE is required for uterine conformational changes in mated Drosophila females. Proc Natl Acad Sci U S A. 106 (37), 15796–15800, (2009).

14 Adams, E. M. & Wolfner, M. F. Seminal proteins but not sperm induce morphological changes in the Drosophila melanogaster female reproductive tract during sperm storage. J Insect Physiol. 53 (4), 319–331, (2007).

15 Singh, A. et al. Long-term interaction between Drosophila sperm and sex peptide is mediated by other seminal proteins that bind only transiently to sperm. Insect Biochem Mol Biol. 102 43–51, (2018).

16 Avila, F. W. & Wolfner, M. F. Cleavage of the Drosophila seminal protein Acp36DE in mated females enhances its sperm storage activity. J Insect Physiol. 101 66–72, (2017).

17 Chapman, T. & Davies, S. J. Functions and analysis of the seminal fluid proteins of male Drosophila melanogaster fruit flies. Peptides. 25 (9), 1477–1490, (2004).

18 Gligorov, D., Sitnik, J. L., Maeda, R. K., Wolfner, M. F. & Karch, F. A novel function for the Hox gene Abd-B in the male accessory gland regulates the long-term female post-mating response in Drosophila. PLoS Genet. 9 (3), e1003395, (2013).

19 Minami, R. et al. The homeodomain protein defective proventriculus is essential for male accessory gland development to enhance fecundity in Drosophila. PLoS One. 7 (3), e32302, (2012).

20 Maeda, R. K. et al. The lncRNA male-specific abdominal plays a critical role in Drosophila accessory gland development and male fertility. PLoS Genet. 14 (7), e1007519, (2018).

21 Prince, E. et al. Rab-mediated trafficking in the secondary cells of Drosophila male accessory glands and its role in fecundity. Traffic. 20 (2), 137–151, (2019).

22 Robinson, M. D. & Oshlack, A. A scaling normalization method for differential expression analysis of RNA-seq data. Genome Biol. 11 (3), R25, (2010).

23 Khan, S. J., Abidi, S. N., Tian, Y., Skinner, A. & Smith-Bolton, R. K. A rapid, gentle and scalable method for dissociation and fluorescent sorting of imaginal disc cells for mRNA sequencing. Fly (Austin). 10 (2), 73–80, (2016).

24 Taniguchi, K. et al. Isoform-specific functions of Mud/NuMA mediate binucleation of Drosophila male accessory gland cells. BMC Dev Biol. 14 46, (2014).

25 Corrigan, L. et al. BMP-regulated exosomes from Drosophila male reproductive glands reprogram female behavior. J Cell Biol. 206 (5), 671–688, (2014).

26 Leiblich, A. et al. Bone morphogenetic protein- and mating-dependent secretory cell growth and migration in the Drosophila accessory gland. Proc Natl Acad Sci U S A. 109 (47), 19292–19297, (2012).

27 Kubo, A. et al. Nutrient conditions sensed by the reproductive organ during development optimize male fecundity in Drosophila. Genes Cells. 23 (7), 557–567, (2018).

